# Genome-Wide Association Study for Yield and Yield related traits reveals Marker–Trait Associations in Germplasm lines of Rice

**DOI:** 10.1101/2023.07.10.548364

**Authors:** Darmagaru Shivani, Abdul Fiyaz Rahaman, Farzana Jabeen, Jukanti Aravind Kumar, Chaithanya Kasarla, Dileep Kumar Gowdru Dhananjaya, Lella Venkata Subba Rao, Supriya, Shoba Venkatanagappa, Raman Meenakshi Sundaram

**Author notes:** Corresponding author: ^1^Department of Plant Breeding, ICAR-Indian Institute of Rice Research, Hyderabad, India, 500030.

## Abstract

Rice germplasm has abundant genetic diversity, which provides a feasible solution for mapping loci of multiple traits simultaneously. In this study, a set of 72 rice germplasm lines were evaluated for yield and yield-related traits, and significant phenotypic variation was observed among the lines. Three accessions with high yield performance were identified. The germplasm set comprised five sub-populations and genome-wide association study (GWAS) identified a total of 6 marker-trait associations (MTAs) for the studied traits. These MTAs were located on rice chromosomes 1, 3, 7, 9, and 12 and explained the trait phenotypic variances ranging from 17.8 to 26.3%. Six novel MTAs were identified for yield and yield-related traits. A total of 28 putative annotated candidate genes were identified in a genomic region spanning ∼200 kb around the MTAs respectively. Among the important genes underlying the novel MTAs were *OsFBK12, bHLH, WRKY, HVA22,* and *ZmEBE-1*, which are known to be associated with the identified novel QTLs. These MTAs provide a pathway for improving high yield in rice genotypes through molecular breeding.

## Introduction

Rice (*Oryza sativa* L.) is the most important agricultural crops and the staple food of various developing countries of the world. In India, rice is cultivated in an area of 4.4 x 107 hm^2^ and provides food for nearly three fourth of the population (Fiyaz et al. 2022). It is a major source of food, also called as ‘Grain of Life’ (Shivani et al. 2021) and provides 21% of the energy and 15% of the protein requirement of human beings (Kennedy 2002). With constant increase in human population, particularly in developing countries where rice is the main source of caloric intake, coupled with climate change and the intensive water, land, and labor requirements of rice cultivation creates a pressing and continuous global need for new, enhanced, resource-use efficient and highly productive rice varieties (Vinci et al. 2023).Currently, breeding for rice faces the problem of yield plateaus, as we witness declining rice productivity in many countries, resulting from the tapered genetic base of parental material used in the breeding programme. Hence, exploiting new QTLs for grain yield is a novel technique for improving rice yield with marker-assisted selection (Kim et al. 2009).

Despite the availability of all the scientific resources, most of what we know about the genetic architecture of complex traits in rice is based on traditional quantitative trait locus (QTL) linkage mapping using bi-parental populations. Due to the presence of limited genetic variation and recombination in the mapping population’s, traditional QTL analysis has restrictions for finding valuable natural variations in trait-associated loci (Chen et al. 2014).

The advancement of techniques and rapid efforts of scientists to obtain significant markers to access trait association have aided the development of less expensive genotyping services also known as genome-wide association studies (Korte and Farlow 2013). Genome-wide association study (GWAS) is an important tool, with enormous potential to accelerate breeding as it enables breeders to make the selection based on marker-trait associations (MTAs) as a response to the combined effect of all favorable alleles. To achieve greater resolution of association analyses, a large number of molecular marker sets are preferred for association mapping. SNPs are the most promising molecular markers used, they are the most abundant form of genetic variation within genomes (Zhu et al. 2008) and a extensive array of technologies have been developed for high-throughput genotyping. Combined, these factors make SNPs ideal source for GWAS using large samples and high-density markers, which greatly improves the resolution of association mapping. Rice germplasm offers a feasible solution of mapping loci of multiple traits simultaneously hence providing a mean to accurately identify the trait expressing genes/SNPs owing to abundant genetic diversity using GWAS. It also enables to simultaneously screen a very huge number of accessions for genetic variation underlying diverse complex traits (McNally et al. 2009).

The present study aims at studying rice germplasm for yield and yield related traits to identify high yielding cultivars and MTAs governing yield and yield related traits. These donors and MTAs can be utilized in breeding programs to develop varieties with high yielding through marker assisted selection.

## Material and Methods

### Plant Materials

A set of 72 rice germplasm lines collected from all over India were evaluated for ten yield and yield related traits. The list of the germplasm lines used in the current study are presented in Supplementary Table 1.

### Evaluation for yield and yield related traits

Seventy two germplasm lines were sown in dry bed during *Kharif* 2019 and *Kharif* 2020 at the ICAR-Indian Institute of Rice Research, Hyderabad. Twenty-seven days old seedlings of each germplasm lines were transplanted by adopting a spacing of 20 cm between rows and 15 cm between plants with two replications in Randomized Block Design. All the necessary precautions were adopted to maintain a uniform plant population per replication. Mean data were collected for ten quantitative characters at the appropriate growth stage of the plant. The characters that were evaluated included days to 50% flowering, plant height (cm), panicle length (cm), number of tillers per plant, number of productive tillers, panicle weight, number of filled grains, number of unfilled grains, 1000 seed weight (g) and grain yield per plant (g). After computing the mean data for each character was subjected to standard methods of analysis of variance following (Panse and Sukhatme 1985).

### Data Analysis

Correlation coefficients were calculated using the formulae suggested by (Falconer 1964) and path analysis by (Dewey and Lu 1959). Hierarchical cluster analysis was conducted using Tocher method using INDOSTAT software.

### DNA Isolation and SNP Genotyping

Total genomic DNA was isolated from young leaves using Cetyl Trimethyl Ammonium Bromide (CTAB) method (Doyle and Doyle 1990). The DNA was quantified using a nano-drop spectrophotometer (NanoDropTM 2000/2000 c, Thermo Fisher Scientific, DE, United States). High throughput genotyping was carried out using Illumina rice 7K Chip, IRRI containing a total of 7,000 high-quality SNPs covering all 12 rice chromosomes. DNA amplification, fragmentation, chip hybridization, single-base extension through DNA ligation, and signal amplification was carried out using the method suggested by (Singh et al. 2015).

### Population Structure Analysis

The software STRUCTURE 2.3.4 (Pritchard et al. 2000) was utilized to determine the population structure of 72 genotypes using a Bayesian model of ADMIXTURE (Alexander et al. 2009), wherein 5 independent runs with 50,000 burn-ins and Markov Chain Monte Carlo (MCMC) period set to 50,000 was conducted for each K. Furthermore, STRUCTURE HARVESTER (Earl and Vonholdt 2011) was used to estimate the optimum number of sub-populations (Evanno et al. 2005).

### Genome-Wide Association Study

Association mapping panel of 72 germplasm lines were genotyped using 7 K SNP chip. SNP data was filtered for minor allele frequency (MAF) 0.05 and maximum missing sites per SNP was fixed to <20%. After filtering, a total of 3,944 SNPs were used to detect MTAs. MTAs were identified using GLM and MLM (Liu 2015, Liu et al. 2016) implemented in GAPIT (genomic association and prediction integrated tool).

## Results

### Phenotypic Evaluation

In the present study, significant phenotypic variation was observed among all the seventy two germplasm lines for all the ten yield and yield related traits under two years of evaluation (Supplementary Table 2, Fig. 1). Days to 50% flowering recorded a general mean value of 100 d ranging from 63 d (Varalu) to 129 (Ketekijoha, Dinesh, Jalmagna) combined over both the seasons. The plant height varied from 81 (Karjat 7) to 156 cm (Dinesh) with the mean value of 107 cm. Incase of panicle length, longest panicle was of the germplasm line Akshyadhan (31.3 cm) while the shortest one was of Vandana (21.4 cm). The mean value for number of tillers per plant was 14 with a range from 10 (Vivekdhan 62) to 18 (Tetep). The mean value of productive tillers per plant was 12 with a range from 9 (Akshyadhan, Aadur-1) to 16 (Taroari basmati, Tetep). With respect to the panicle weight, more panicle weight was reported for germplasm line NDR 359 (6.05 g) while the lowest one was of Sukaradhan-1 (1.5 g). A maximum of 223 and a minimum of 53 filled grains per panicle were observed in germplasm lines Pushyami and Sukaradhan-1, respectively. The 1000 seed weight ranged from 12.1 g (Chittimutyalu) to 28.95 g (Pantdhan 19). Grain yield per plant ranged from 8.6 g (Taraori basmati) to 24.45 g (Co 50) with a general mean of 16 g. The genetic variability is depicted in the form of box plots (Fig. 1) showed the frequency distribution for ten quantitative traits among 72 germplasm lines of O. sativa. The traits, namely, number of filled grains per panicle, number of unfilled grains per panicle and panicle weight, exhibited greater genetic variability in both the seasons *Kharif* 2019 and *Kharif* 2020 (Supplementary Table 3).

**Fig. 1.**
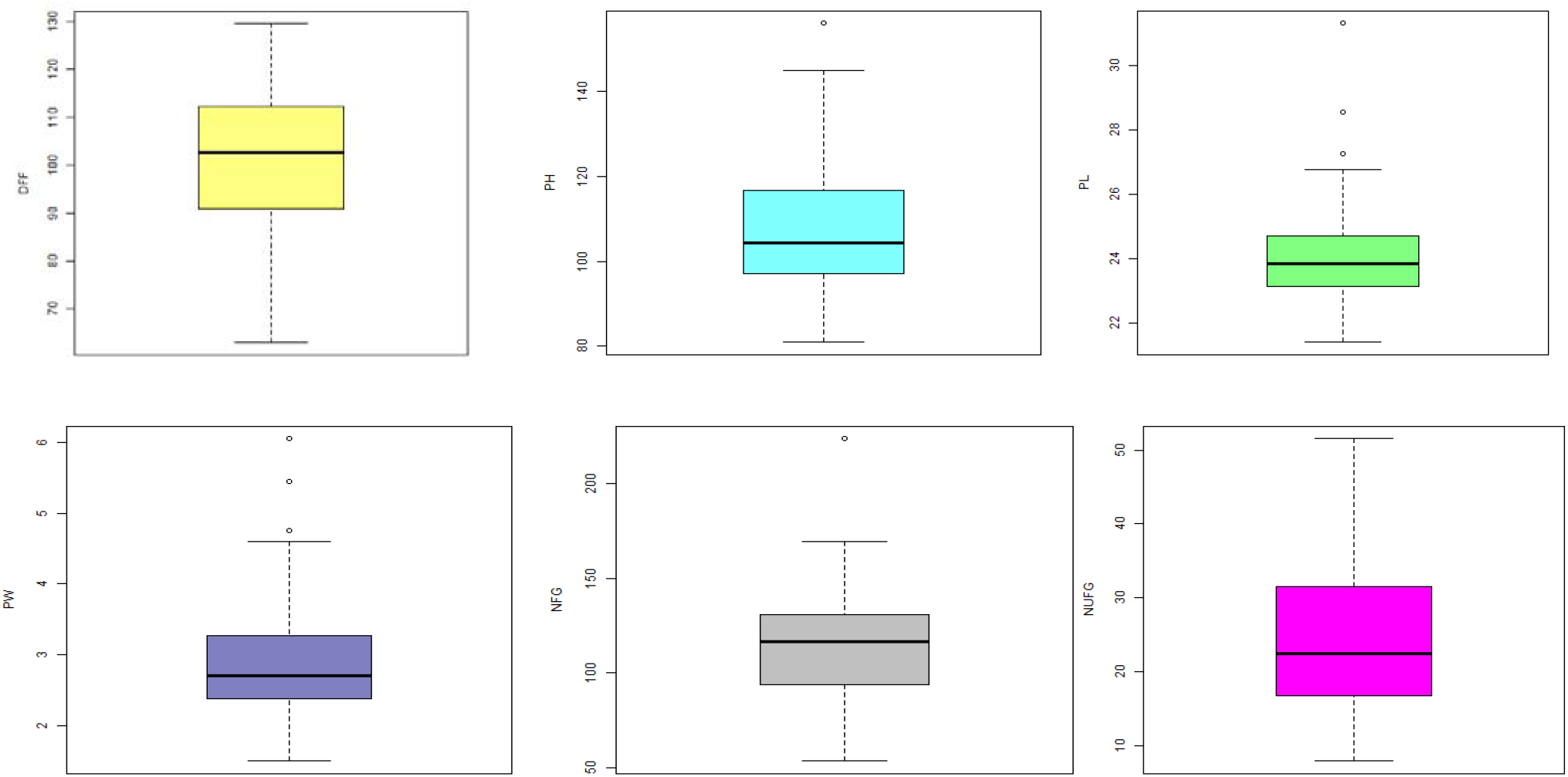
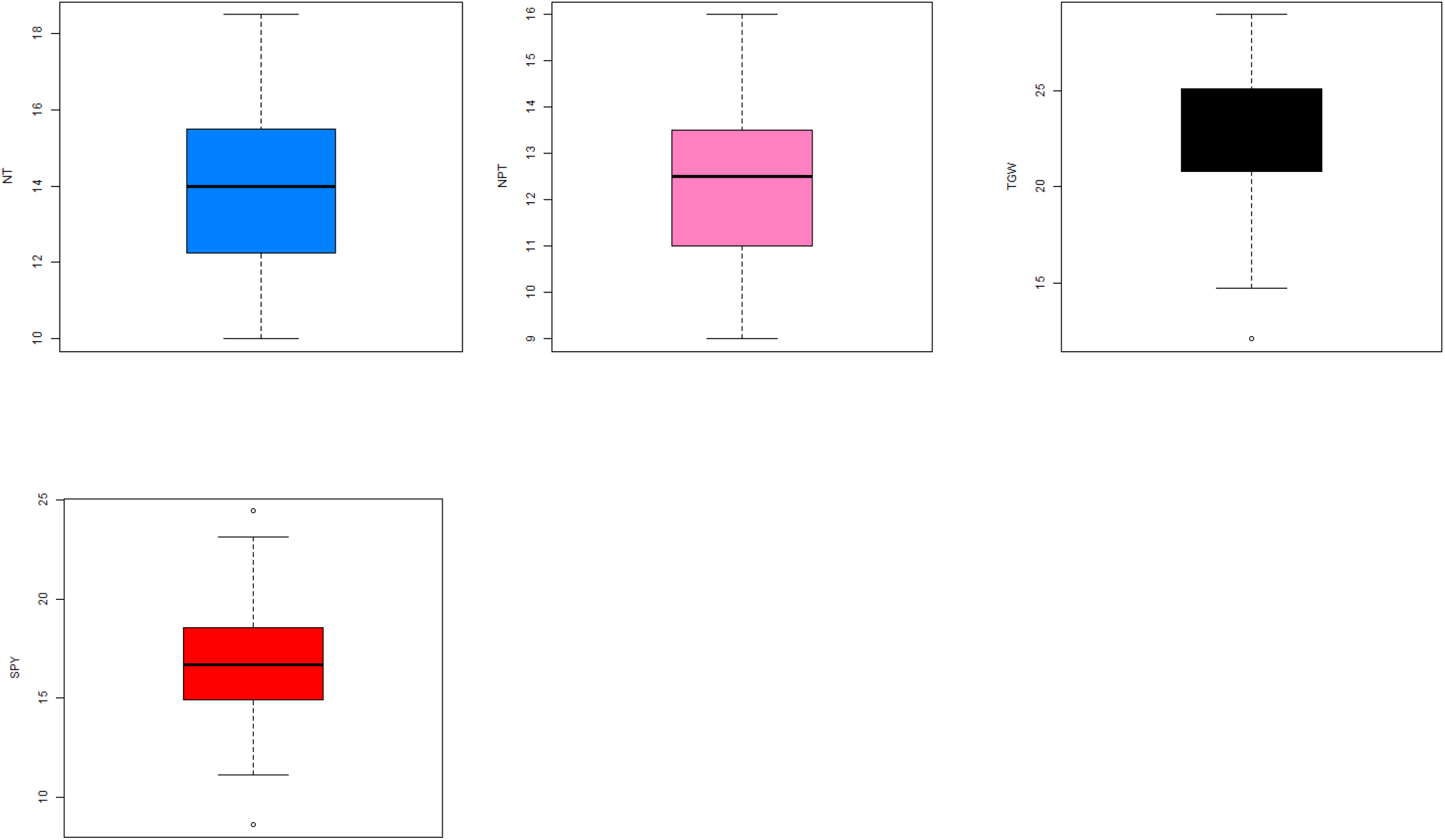
Box-plots showing the variation of the data from the ten yield variables of two seasons evaluated in 72 germplasm lines of rice. The upper, median and lower quartiles represent the 75th, 50th and 25th percentiles of the germplasm respectively. The vertical lines represent the variation in the population. Dots represent the outliners.

A dendrogram was constructed on the basis of yield and yield related traits to classify the germplasm lines. In the present study, heatmap (Supplementary Fig. 1) revealed number of filled grains per panicle exhibiting greater variation among the accessions followed by plant height and days to 50% flowering. The 72 germplasm lines were grouped into nine clusters based on D2 values using the Tocher method. The distribution of germplasm lines into various clusters is displayed in (Supplementary Table 4, Supplementary Fig. 2). Out of the nine clusters, cluster I was the largest comprising of 24 germplasm lines followed by cluster II with twenty one, cluster IV and cluster III with 13 and 7 genotypes each. Clusters V, VI, VIIII and IX had only one genotype each. The cluster means for each of 10 characters (Supplementary Table 5) indicated that cluster IX had many of the desirable means for most of the characters and with respect to relative contribution of different characters to the genetic diversity (Supplementary Table 6), the contribution of days to 50% flowering was highest (70.8%) followed by plant height (16.16%), number of filled grains per panicle (4.73%), single plant yield (3.71%), 1000 grain weight (3.56%), panicle weight (0.66), number of unfilled grains per panicle (0.15), panicle length (0.11), number of productive tillers per plant (0.09) and total number of tillers per plant (0.03). The characters days to 50% flowering and plant height contributed 86.96% towards total divergence.

### Correlation and path analysis among yield and yield related traits

Correlation analysis among yield and its attributing traits depicted in the form of correlogram (Fig. 2) revealed that grain yield per plant had significant positive correlation with days to 50% flowering (0.16), panicle weight (0.24) and number of filled grains per panicle (0.23), while a significant negative association was found with total number of tillers per plant (−0.12). Whereas, path coefficient analysis revealed the trait, number of productive tillers exerting the highest direct positive effect (0.824) on grain yield per plant followed by panicle length (0.193), number of filled grains per panicle (0.102), days to 50% flowering (0.10), panicle weight (0.099) and 1000 grain weight (Supplementary Table 7). While, negative direct effect on grain yield was recorded by plant height (−0.163) and total number of tillers per plant (−0.915). The high positive indirect effects on grain yield per plant was exhibited by total number of tillers per plant via panicle length (0.205), panicle weight (0.165), number of filled grains per panicle (0.143), thousand grain weight (0.135) and plant height (0.025).

**Fig. 2.**
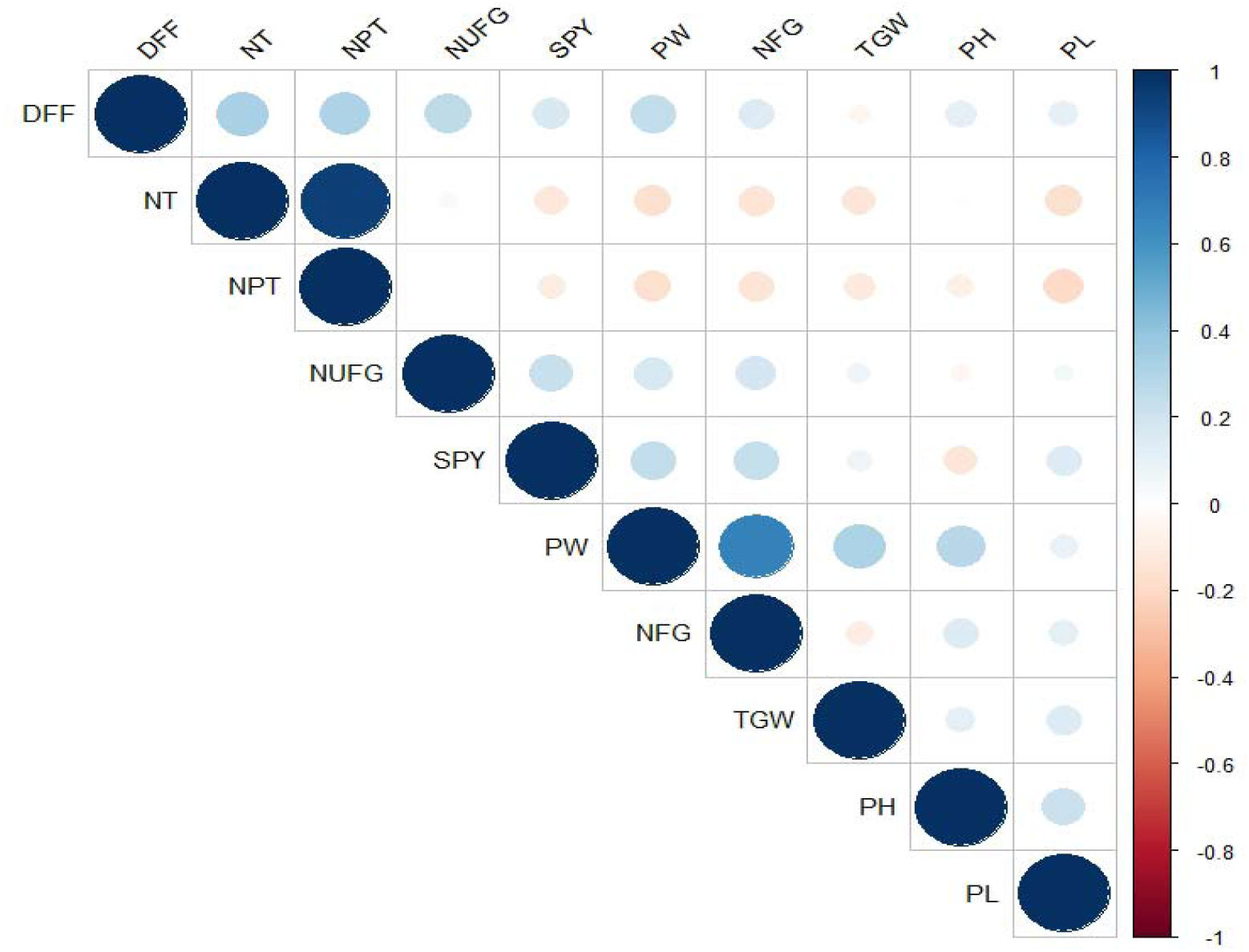
Correlation coefficients among various yield and yield related traits. **DFF- Days to 50% flowering, PH- Plant height, PL- Panicle length, TNT- Total number of tillers, PNT- Productive tillers, PW- Panicle weight, NFG- Number of filled grains, NUFG- Number of unfilled grains, TGW- Thousand grain weight, SPY- Single plant yield**

### Population Structure

A set of 72 germplasm lines in the current study was subjected to population structure analysis. Based on Evanno plot, ΔK value was highest for the model parameter K=5 (Fig. 3a). Therefore, the optimal number of sub-populations (K) was determined to be 5, which are represented as POP1, POP2, POP3, POP4 and POP5 (Fig. 3b). The fixation index (Fst) was 0.95, 0.8, 0.79, 0.9 and 0.9 for the sub-populations POP1, POP2, POP3, POP4 and POP5 respectively. POP1 was the largest sub-population and constituted 28 genotypes, POP2 comprised 12 genotypes, POP3 comprised 16, POP4 and POP5 comprised 11 and 5 genotypes indicating sufficient amount of diversity existing in the germplasm lines studied.

**Fig. 3.**
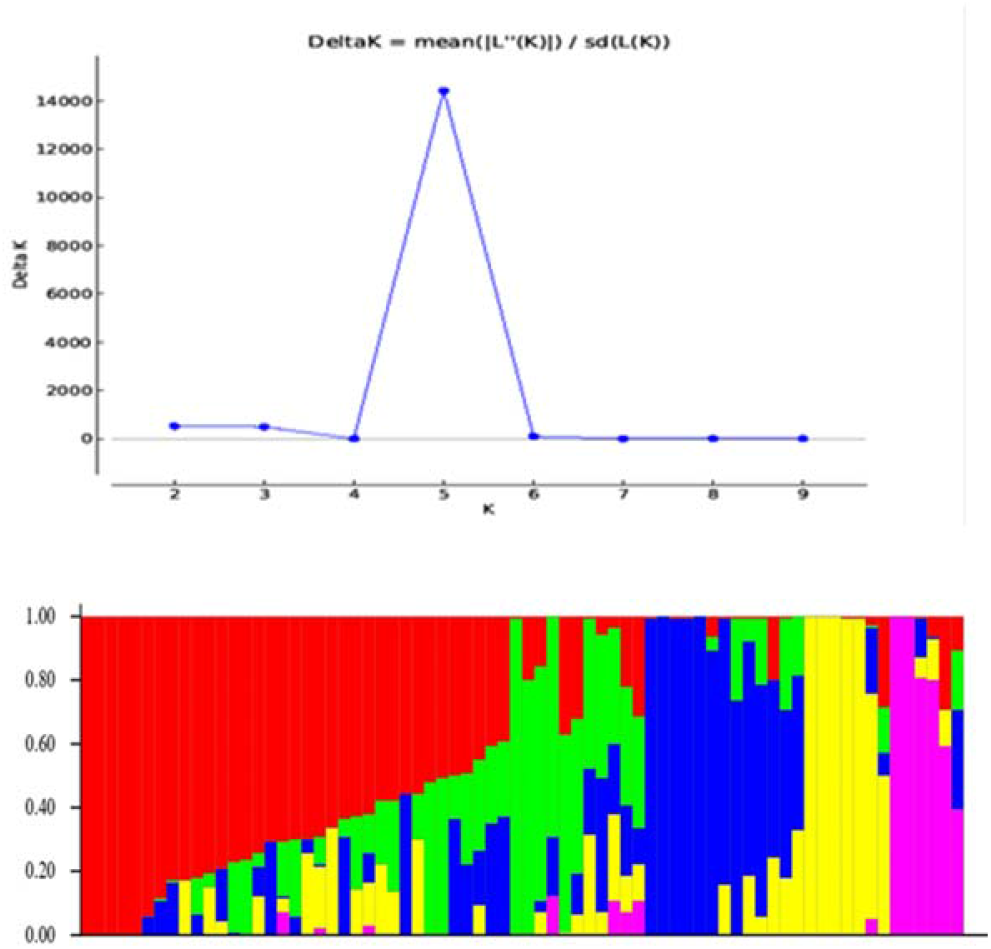
STRUCTURE analysis. (**a**)The ΔK value was highest for the model parameter K=3than for other values of K. (**b**)The bar plot showing each rice variety belonging to three subpopulations

### Genome-Wide Association Study for Traits Associated with Salinity Tolerance

A total of 6novel MTAs were identified for ten yield and yield related traits in the current study. These MTAs are located on rice chromosomes 1, 3, 7, 9and 12 and explained the trait phenotypic variance ranging from 17.8% to 26.3% (Table 1). Manhattan plots and quantile–quantile (Q-Q) plots generated through the model are presented in Fig. 4. The Q-Q plots indicate that the model was well fitted to the data as the observed p-values showed less deviation from the expected p- values.

**Fig. 4.**
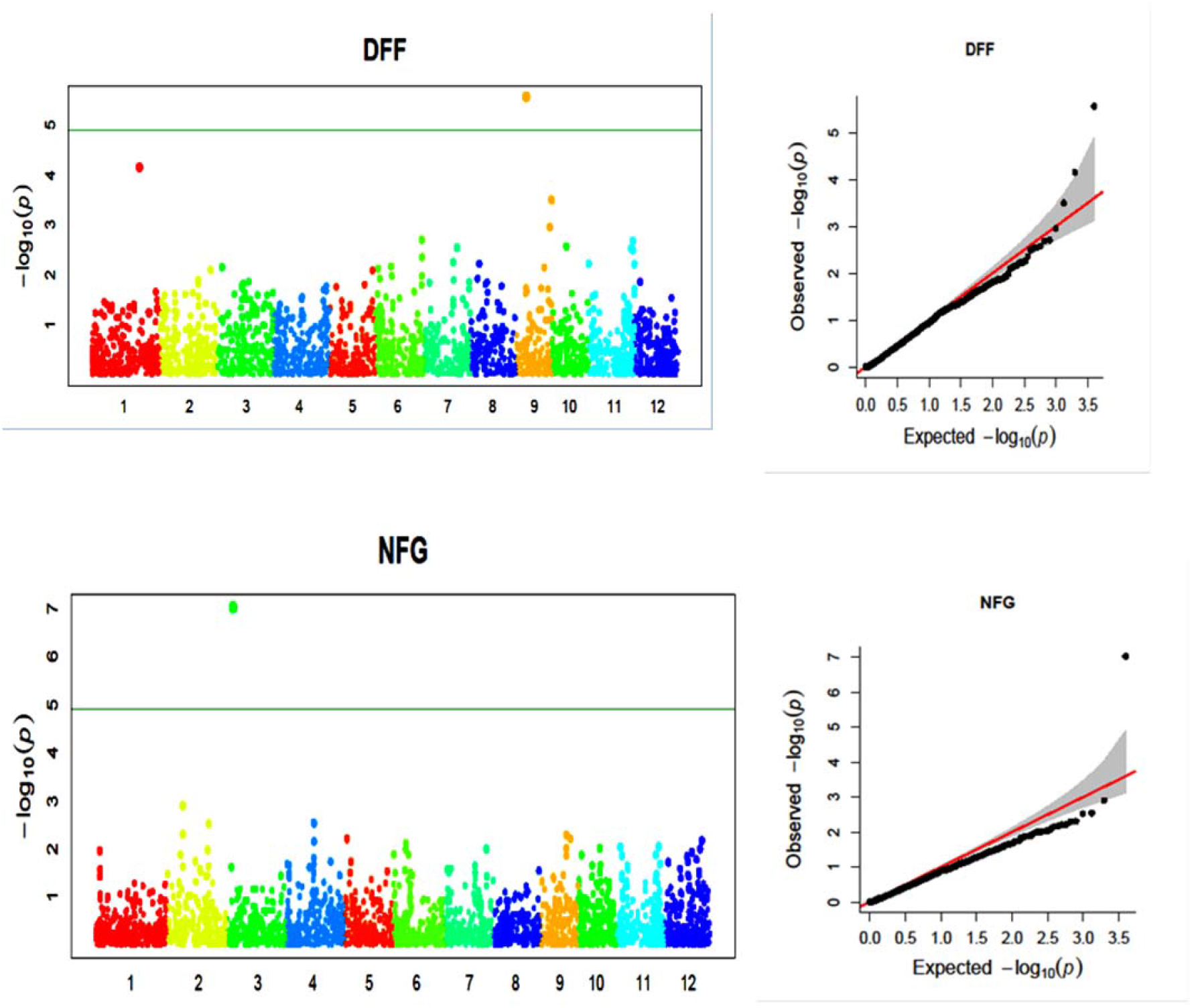
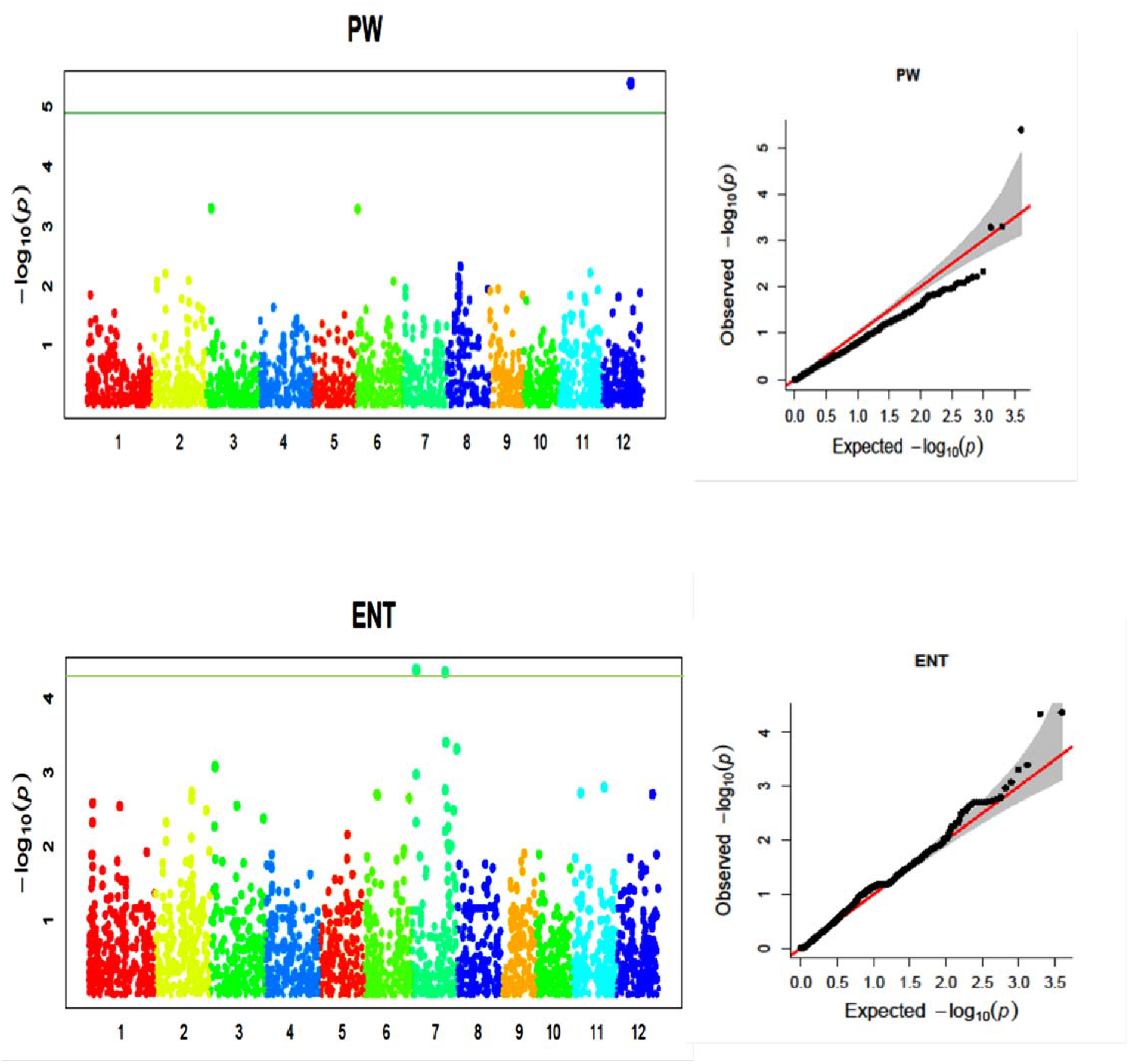
Manhattan plots and Q–Q plots representing the significant marker trait associations for 4 yield related traits. DFF, Days to 50% flowering, NFG, Number of filled grains per panicle, PW, Panicle weight (g), ENT, Effective number of tillers.

**Table 1.** Details of the marker-trait associations(MTAs)identified for yield and yield related traits.

| Character |  | Marker | Chr. | Position (bp) | P-value | Marker R<sup>2</sup> | Previous report |
| --- | --- | --- | --- | --- | --- | --- | --- |
| **DFF** | qDFF1 | S_1007944 | 1 | 30373171 | 0.0007971 | 24.9 | - |
|  | qDFF9 | S_9311693 | 9 | 6360984 | 0.002143 | 24.9 | - |
| **PW** | qPW12 | S_12807606 | 12 | 19376763 | 0.000574 | 17.8 | - |
| **NFG** | qNFG3 | SNP-3.3850394 | 3 | 3851399 | 5.28E-05 | 26.3 | - |
| **ENT** | qENT7 | S_7032438 | 7 | 2679227 | 4.25E-05 | 22.6 | - |
|  | qENT7.1 | SNP-7.21505364 | 7 | 21506358 | 4.55E-05 | 22 | - |

For DFF, two novel QTLs, S_1007944, S_9311693 were identified on chromosome 1 and 9 and explained the phenotypic variance of 24.9%. For PW, one novel QTL (S_12807606) was identified on chromosome 12 with phenotypic variance of 17.8%.

One novel QTL (SNP-3.3850394) was identified for number of filled grains per panicle on chromosome 3 with variance of 26.3%. For effective number of tillers per plant two novel QTLs (S_7032438 and SNP-7.21505364) were identified on chromosome 7 with phenotypic variance of 22 and 22.6%, respectively.

*Insilico* search for annotated gene MSU-RAP database led to identification of 28putative candidate genes in ∼200 Kb genomic region of identified novel MTAs respectively (Table 2).

**Table 2.** Putative candidate genes annotated in ∼200 Kb genomic regions in identified novel MTAs.

| Characters | Chr. | Gene | Position | Function |
| --- | --- | --- | --- | --- |
| **DFF** | 1 | LOC_Os01g52780 | 30370212 | HVA22, putative, expressed |
|  |  | LOC_Os01g52790 | 30376917 | cytochrome P450 72A1, putative, expressed |
|  |  | LOC_Os01g52830 | 30387917 | DUF1264 domain containing protein, putative, expressed |
|  |  | LOC_Os01g52880 | 30407036 | leucine-rich repeat family protein, putative, expressed |
|  |  | LOC_Os01g52970 | 30449206 | OsFBX22 - F-box domain containing protein, expressed |
|  |  | LOC_Os01g53070 | 30496722 | ATP binding protein, putative, expressed |
|  | 9 | LOC_Os09g11450 | 6383644 | transporter, monovalent cation:proton antiporter-2 family, putative, expressed |
|  |  | LOC_Os09g11460 | 6389789 | AP2 domain containing protein, expressed |
|  |  | LOC_Os09g11510 | 6438925 | phosphoglyceratemutase, putative, expressed |
|  |  | LOC_Os09g11520 | 6441905 | DUF565 domain containing protein, putative, expressed |
| **ENT** | 7 | LOC_Os07g05650 | 2690658 | MAGE-8 antigen-related, putative, expressed |
|  |  | LOC_Os07g05640 | 2686437 | Transporter family protein, putative, expressed |
|  |  | LOC_Os07g05720 | 2751213 | TCP family transcription factor, putative, expressed |
|  |  | LOC_Os07g05800 | 2791450 | Glutathione S-transferase, N-terminal domain containing protein, expressed |
|  |  | LOC_Os07g35970 | 21519953 | SMP-30/Gluconolactonase/LRE-like region containing protein, expressed |
|  |  | LOC_Os07g35940 | 21506356 | beta-amylase, putative, expressed |
| **PW** | 12 | LOC_Os05g03810 | 19381453 | Trehalose phosphatase, putative |
|  |  | LOC_Os12g32040 | 19333950 | ZmEBE-1 protein, putative, expressed |
|  |  | LOC_Os12g32180 | 19424812 | cornichon protein, putative, expressed |
|  |  | LOC_Os12g32210 | 19441176 | autophagy-related protein 10, putative, expressed |
|  |  | LOC_Os02g15870 | 19452169 | molybdenum cofactor biosynthesis protein 1, putative, expressed |
|  |  | LOC_Os12g32250 | 19473728 | WRKY DNA-binding domain containing protein, expressed |
| **NFG** | 3 | LOC_Os03g07580 | 3859385 | OsNucAP1 - Putative NucleoporinAutopeptidase homologue, expressed |
|  |  | LOC_Os03g07600.1 | 3871355 | Calcium-binding protein, putative, expressed |
|  |  | LOC_Os03g07790 | 3960943 | Zinc finger, C3HC4 type domain containing protein, expressed |
|  |  | LOC_Os03g07450 | 3786933 | homeobox associated leucine zipper, putative, expressed |
|  |  | LOC_Os03g07530 | 3833954 | OsFBK12 - F-box domain and kelch repeat containing protein, expressed |
|  |  | LOC_Os10g26460 | 3845713 | bHLH family protein, putative, expressed |

## Discussion

Rice (*Oryza sativa* L.) is one of the most important agricultural crops and the main staple food of various developing countries. Rice yield is one of the most important and complex traits governed by polygenes. In rice, the yield is determined by indirect traits like plant height, growth period, tillering ability, panicle length, seed length, seed setting rate, and grains per panicle, as well as direct traits like panicle number per unit area and/or per plant, filled grains per panicle and 1000-grain-weight. With the constant increase of the world population and the decline of land resources, we are in great requisite to continuously improve the grain yield of rice. Therefore, development of rice varieties with high yield is of utmost importance. In the present study, we have evaluated a set of germplasm lines for yield and yield related traits. We observed the tremendous diversity for all measured traits and identified two high yielding lines (Co 50, Akshyadhan and Pushyami) and having positive alleles for yield and yield related traits. These lines can be used as donor source in breeding programs for development of high yielding rice varieties.

Correlation studies help in examining the possibility of improving yield through indirect selection of its component traits which are highly correlated. Since grain yield per plant was positively correlated with days to 50% flowering, panicle weight and number of filled grains per panicle, direct selection for these traits could be considered as the criteria for higher grain yield as reported by (Sarker et al. 2014) for days to 50% flowering, number of filled grains per panicle and (Prasad et al. 2017) for panicle weight. The total number of tillers per plant showed significant negative correlated with yield per plant. Thus, this trait may not be rewarding if selected for enhancing the grain yield. Path analysis revealed the traits, number of productive tillers, panicle length and number of filled grains per panicle exerted the highest direct positive effect on grain yield per plant indicating that selection for these characters is likely to bring about an overall improvement in grain yield.

Identifying the genomic regions governing this complex trait is of utmost importance to develop high yielding rice varieties. Genome-wide association study (GWAS) is an important tool, with enormous potential to accelerate breeding as it enables breeders to make the selection based on marker-trait associations (MTAs) as a response to the combined effect of all favorable alleles.

In the current study, five sub-populations were detected which were considered in mixed linear model (MLM) to reduce spurious associations. MLM has been popularly adopted for GWAS in crops (Zhang et al. 2014, Ya-Fang et al. 2015, Wang et al. 2016).The mixed linear model uses both fixed and random effects which incorporates kinship among the individuals. Though, MLM being a single locus method that allows testing of one marker locus at a time, had an intrinsic limitation in matching the real genetic architecture of the complex traits that are under the effect of multiple loci acting simultaneously (Kaler and Purcell 2019). Population structure and individuals’ total genetic effects are often fitted as covariates in a mixed linear model (MLM) to reduce the false discovery rate (FDR) (Yu et al. 2006).

The GAPIT implemented several new methods that improve statistical power and computing speed for data analyses (Fig. 4). Furthermore, the GAPIT includes an enhanced set of data interpretation and evaluation functions.The threshold of -log10(P) >3 was used to declare MTAs because of restricted number of genotypes used in the study. In one of the recent studies, (Yadav et al. 2021) conducted GWAS for salinity tolerance in 96 accessions of rice and found two significant SNPs. (Rohilla et al. 2020) used 94 deep-water rice genotypes of India in GWAS for anaerobic germination (AG) and found significant associated SNPs at log10(P) =3. Similarly, (Feng et al. 2016) performed GWAS for grain shape traits in indica rice and found significant associated SNPs at log10(P) = 3. (Jha et al. 2019) found significant SNPs for agronomic and seed quality traits of pea at log10(P) =3.

In the present study, we have identified a total of 6 MTAs for yield and yield related traits. These QTLs are located on rice chromosomes 1, 3, 7,9 and 12 and explained the trait phenotypic variances ranging from 17.8% to 26.3%. We compared the MTAs identified in this study with previously reported QTLs related to yield and yield related traits in the QTL Annotation Rice Online (Q-TARO) database and by literature survey. This comparison showed that none of the MTA identified in this study was located either in or near the previously reported QTLs associated with yield and yield related traits (Table 1).

In the present study we detected 6 novel MTAs for days to fifty percent flowering, panicle weight, effective number of tillers per plant and number of filled grains per panicle. In silico analysis revealed that the candidate gene LOC_Os01g52780 in the genomic region of MTA qDFF1, was annotated as HVA22 (Table 2). HVA22 is an ER- and Golgi-localized protein capable of negatively regulating GA-mediated vacuolation or programmed cell death in aleurone cells. ABA will induce the accumulation of HVA22 proteins to inhibit vesicular trafficking involved in nutrient mobilization to delay coalescence of protein storage vacuoles as part of its role in regulating seed germination and seedling growth (Guo and David 2008). The other putative candidate gene, LOC_Os01g52830 in the genomic regions of MTA qDFF1, was annotated as DUF1264. This protein expresses under response to abscisic acid stimulus and is located in cellular component and expressed in seed; during seedling growth and seed development stages (Xing et al. 2015).

The putative candidate gene, LOC_Os01g18440 in the genomic regions of MTAs qPL1, was annotated as OsMADS89 (MADS-box family gene with M-gamma type-box) respectively (Table 2). MADS-box proteins are expressed in leaves, shoot, root, panicle and seed, involved in variety of plant developmental process, floral induction, development, formation of seed which results in increased number of grains per panicle (Shah et al. 2022).

The putative candidate gene, LOC_Os03g07790 in the genomic region of MTA, qNFG3 was annotated as zinc finger, C_3_HC_4_ type domain containing protein, expressed (Table 2). Zinc finger type proteins found to play an important role in plant architecture and grain yield (Zhou et al. 2017). The putative candidate gene LOC_Os12g32250, in the genomic region of MTA, qPW12was annotated as WRKY DNA-binding domain containing protein. WRKY genes found to play a variety of developmental and physiological roles in plants. The most reported studies for this superfamily of genes address their involvement in salicylic acid (SA) and disease responses. Some *WRKY* genes regulate embryogenesis, seed coat and trichome development and senescence *WRKY* genes may control seed germination and post germination growth as well (Xie et al. 2005). The other putative candidate gene, LOC_Os12g32040 in the genomic regions of MTA qPW12, was annotated as ZmEBE-1 protein. This protein is potentially involved in the early development of specialized domains of the endosperm (Magnard et al. 2003).

The putative candidate gene, LOC_Os10g26460 in the genomic regions of MTAqNFG3, was annotated as bHLH family protein. bHLH family protein shows similar expression patterns in the flower and seed, supporting the hypothesis that these genes function in rice and Arabidopsis reproductive development. Eleven rice bHLH genes have been characterized. *LAX* (*OsbHLH122*) regulates shoot branching, *Udt1* (*OsbHLH164*) is critical for tapetum development, *OSB1* (*OsbHLH013*) and *OSB2* (*OsbHLH016*) are involved in anthocyanin biosynthesis. bHLH genes found to perform a variety of functions in different tissues at multiple developmental stages (Li et al. 2006). The other putative candidate gene, LOC_Os07g35940 in the genomic region of MTA qENT7.1, was annotated as beta-amylase. This gene was located at 21506356bp, which is very close to the location of qtl i.e 21506358bp.

These novel MTAs identified in the study may play important role in improving yield in rice.

## Conflicts of Interest

The authors declare no conflict of interest.

---

**Supplementary Table 1.**
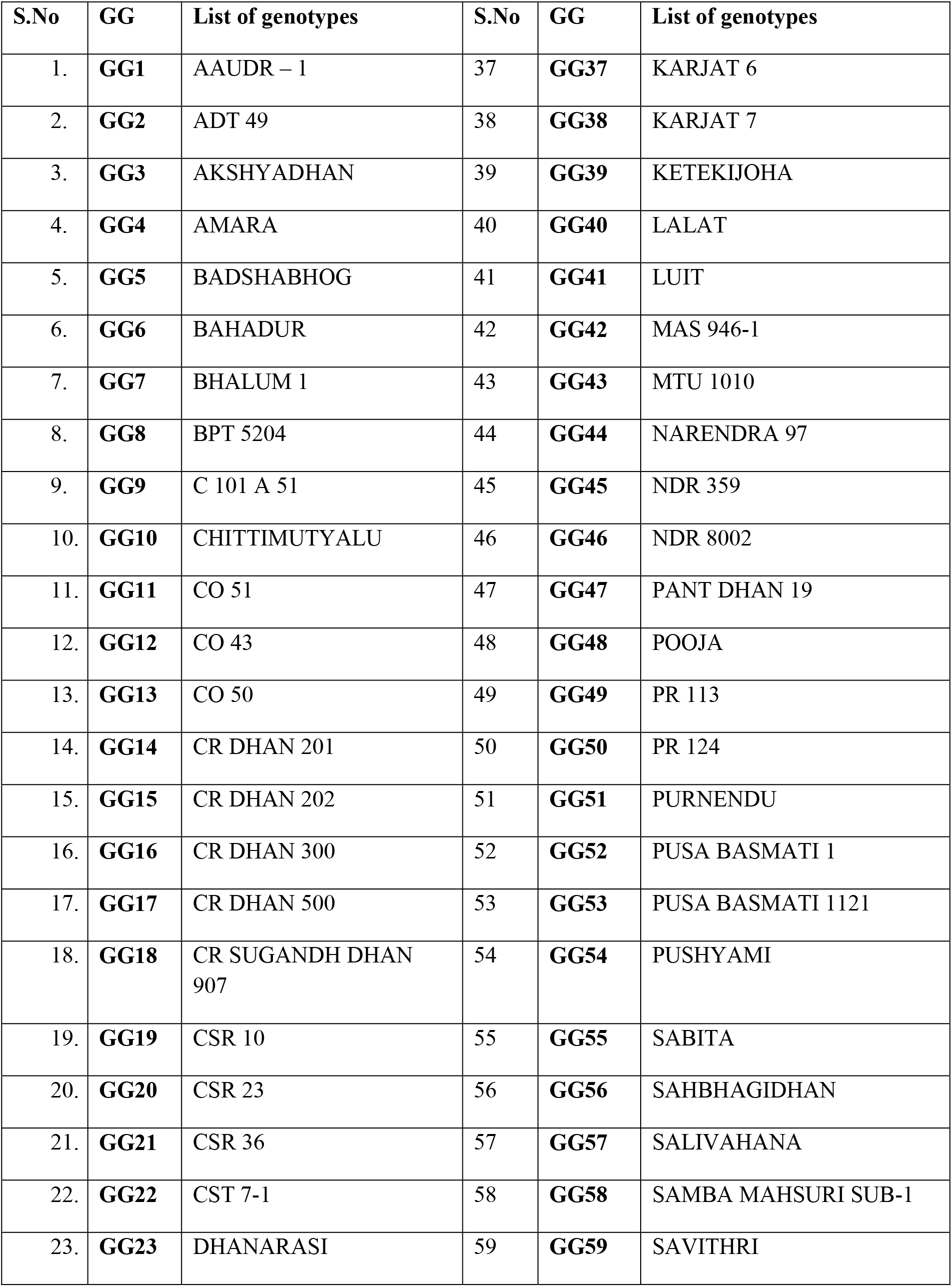
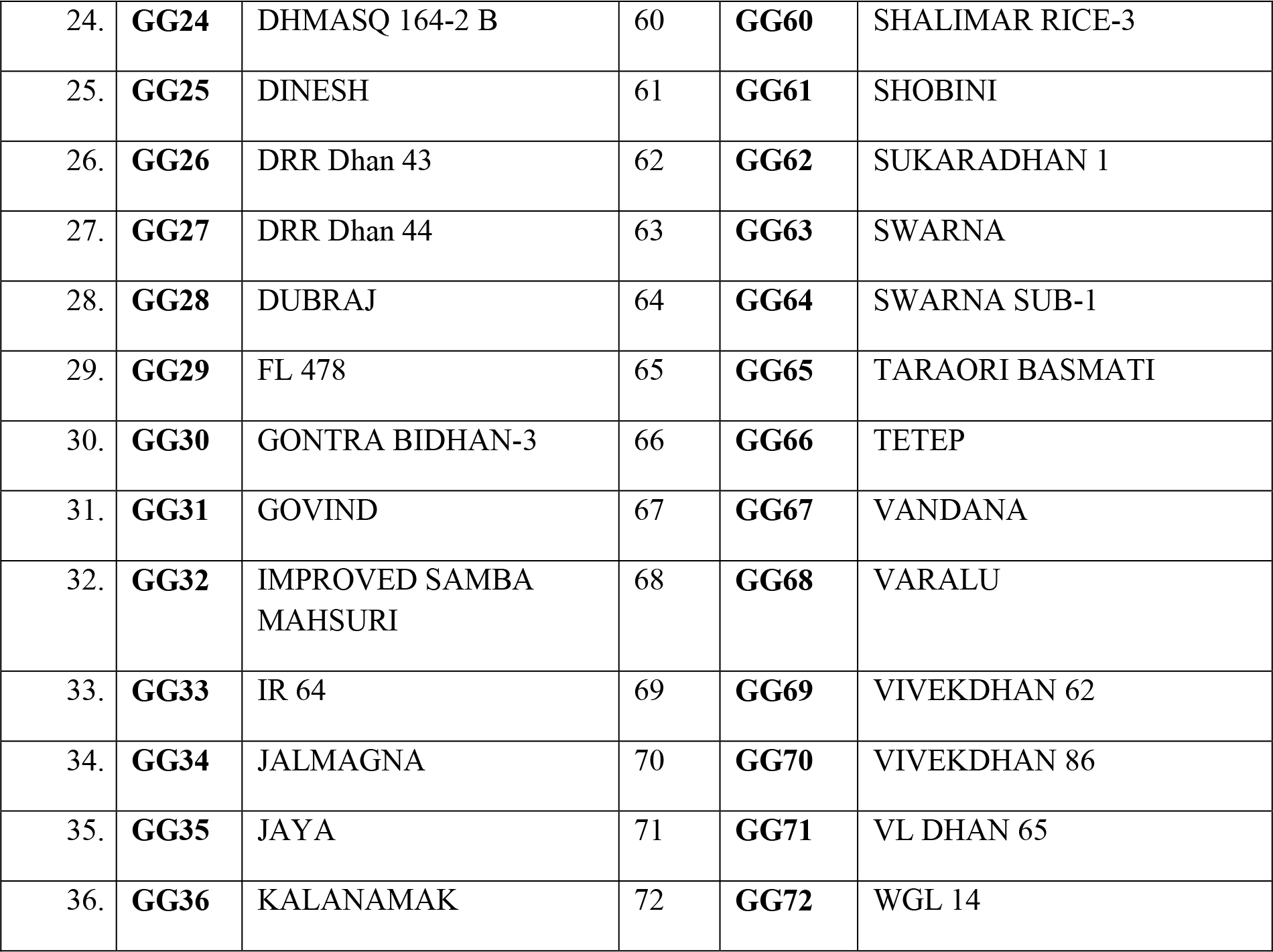
List of germplasm lines used in the present study.

| S.No | GG | List of genotypes | S.No | GG | List of genotypes |
| --- | --- | --- | --- | --- | --- |
| 1. | **GG1** | AAUDR – 1 | 37 | **GG37** | KARJAT 6 |
| 2. | **GG2** | ADT 49 | 38 | **GG38** | KARJAT 7 |
| 3. | **GG3** | AKSHYADHAN | 39 | **GG39** | KETEKIJOHA |
| 4. | **GG4** | AMARA | 40 | **GG40** | LALAT |
| 5. | **GG5** | BADSHABHOG | 41 | **GG41** | LUIT |
| 6. | **GG6** | BAHADUR | 42 | **GG42** | MAS 946-1 |
| 7. | **GG7** | BHALUM 1 | 43 | **GG43** | MTU 1010 |
| 8. | **GG8** | BPT 5204 | 44 | **GG44** | NARENDRA 97 |
| 9. | **GG9** | C 101 A 51 | 45 | **GG45** | NDR 359 |
| 10. | **GG10** | CHITTIMUTYALU | 46 | **GG46** | NDR 8002 |
| 11. | **GG11** | CO 51 | 47 | **GG47** | PANT DHAN 19 |
| 12. | **GG12** | CO 43 | 48 | **GG48** | POOJA |
| 13. | **GG13** | CO 50 | 49 | **GG49** | PR 113 |
| 14. | **GG14** | CR DHAN 201 | 50 | **GG50** | PR 124 |
| 15. | **GG15** | CR DHAN 202 | 51 | **GG51** | PURNENDU |
| 16. | **GG16** | CR DHAN 300 | 52 | **GG52** | PUSA BASMATI 1 |
| 17. | **GG17** | CR DHAN 500 | 53 | **GG53** | PUSA BASMATI 1121 |
| 18. | **GG18** | CR SUGANDH DHAN<br>907 | 54 | **GG54** | PUSHYAMI |
| 19. | **GG19** | CSR 10 | 55 | **GG55** | SABITA |
| 20. | **GG20** | CSR 23 | 56 | **GG56** | SAHBHAGIDHAN |
| 21. | **GG21** | CSR 36 | 57 | **GG57** | SALIVAHANA |
| 22. | **GG22** | CST 7-1 | 58 | **GG58** | SAMBA MAHSURI SUB-1 |
| 23. | **GG23** | DHANARASI | 59 | **GG59** | SAVITHRI |
| 24. | **GG24** | DHMASQ 164-2 B | 60 | **GG60** | SHALIMAR RICE-3 |
| 25. | **GG25** | DINESH | 61 | **GG61** | SHOBINI |
| 26. | **GG26** | DRR Dhan 43 | 62 | **GG62** | SUKARADHAN 1 |
| 27. | **GG27** | DRR Dhan 44 | 63 | **GG63** | SWARNA |
| 28. | **GG28** | DUBRAJ | 64 | **GG64** | SWARNA SUB-1 |
| 29. | **GG29** | FL 478 | 65 | **GG65** | TARAORI BASMATI |
| 30. | **GG30** | GONTRA BIDHAN-3 | 66 | **GG66** | TETEP |
| 31. | **GG31** | GOVIND | 67 | **GG67** | VANDANA |
| 32. | **GG32** | IMPROVED SAMBA MAHSURI | 68 | **GG68** | VARALU |
| 33. | **GG33** | IR 64 | 69 | **GG69** | VIVEKDHAN 62 |
| 34. | **GG34** | JALMAGNA | 70 | **GG70** | VIVEKDHAN 86 |
| 35. | **GG35** | JAYA | 71 | **GG71** | VL DHAN 65 |
| 36. | **GG36** | KALANAMAK | 72 | **GG72** | WGL 14 |

**Supplementary Table 2.** Analysis of variance (Mean sum of squares) for yield and its component traits of two seasons in germplasm lines of rice.

| S. No | Characters | Seasons | Mean Sum of Squares |  |  |
| --- | --- | --- | --- | --- | --- |
|  |  |  | Replications<br>(d.f. = 1) | Treatments<br>(d.f. = 71 ) | Error<br>(d.f. = 71 ) |
| **1** | **Days to 50 percent flowering** | *Kharif2019* | 0.0278 | 470.00** | 0.5630 |
|  |  | *Kharif2020* | 2.3767 | 476.18** | 0.3662 |
| **2** | **Plant height (cm)** | *Kharif2019* | 0.2556 | 455.81** | 1.5523 |
|  |  | *Kharif2020* | 0.2350 | 500.96** | 1.2208 |
| **3** | **Panicle length (cm)** | *Kharif2019* | 0.8050 | 4.62** | 0.3569 |
|  |  | *Kharif2020* | 1.6792 | 5.61** | 0.4124 |
| **4** | **Number of tillers** **per plant** | *Kharif2019* | 0.0934 | 11.28** | 0.7467 |
|  |  | *Kharif2020* | 14.9941 | 7.79** | 0.5497 |
| **5** | **Number of productive tillers per plant** | *Kharif2019* | 0.5216 | 9.43** | 0.6515 |
|  |  | *Kharif2020* | 29.6420 | 6.27** | 0.7453 |
| **6** | **Number of filled grains per panicle** | *Kharif2019* | 32.8234 | 1383.38** | 3.5515 |
|  |  | *Kharif2020* | 70.8403 | 1529.37** | 3.0488 |
| **7** | **Number of unfilled grains per panicle** | *Kharif2019* | 0.6534 | 240.08** | 1.3983 |
|  |  | *Kharif2020* | 5.5748 | 319.79** | 6.4669 |
| **8** | **Panicle weight (g)** | *Kharif2019* | 0.0621 | 1.31** | 0.0448 |
|  |  | *Kharif2020* | 0.0176 | 1.93** | 0.0701 |
| **9** | **1000 grain weight (g)** | *Kharif2019* | 2.5407 | 23.45** | 0.7309 |
|  |  | *Kharif2020* | 0.1907 | 20.34** | 0.2650 |
| **10** | **Grain yield per plant (g)** | *Kharif2019* | 4.1384 | 19.92** | 0.5146 |
|  |  | *Kharif2020* | 3.2716 | 19.88** | 0.7973 |
| ***Significant at 0.1% level, ** Significant at 1% level, * Significant at 5% level |  |  |  |  |  |

**Supplementary Table 3.**
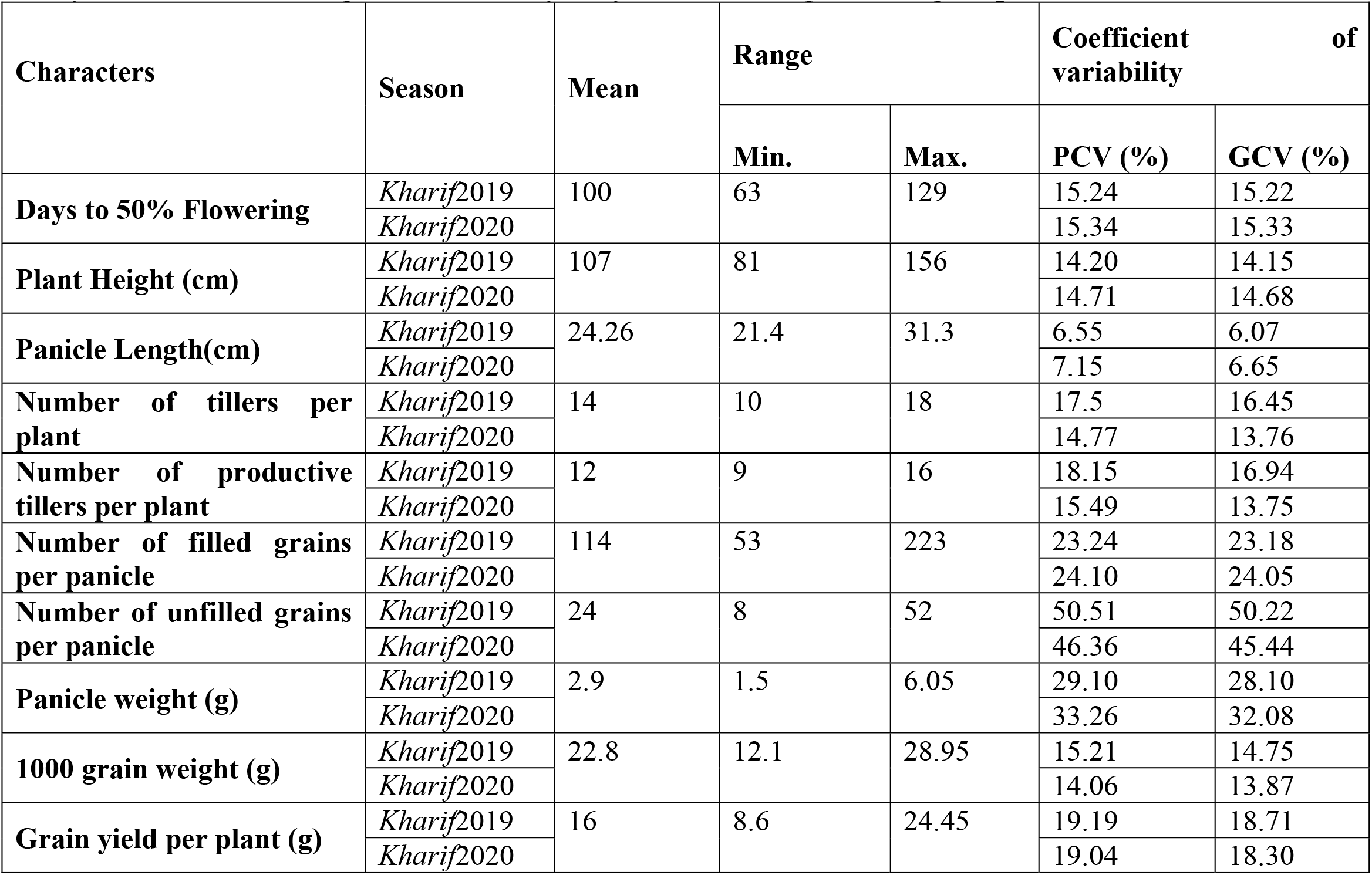
Mean, Range and variability for yield attributing traits in germplasm lines of rice.

**Supplementary Table 4.** Average Inter and intra cluster distances of germplasm lines obtained by D² analysis using ten yield and yield contributing traits.

|  | Cluster 1 | Cluster 2 | Cluster 3 | Cluster 4 | Cluster 5 | Cluster 6 | Cluster 7 | Cluster 8 | Cluster 9 |
| --- | --- | --- | --- | --- | --- | --- | --- | --- | --- |
| Cluster 1 | **188.61** | **539.79** | 2173.3 | 616.5 | 1172.8 | 342.5 | 648.6 | 489.5 | 1820.1 |
| Cluster 2 |  | **231.96** | 904.8 | 1721.0 | 420.6 | 373.3 | 747.2 | 988.9 | 3342.4 |
| Cluster 3 |  |  | **258.75** | 4427.61 | 514.1 | 1554.2 | 2124.8 | 2872.5 | 6787.6 |
| Cluster 4 |  |  |  | **203.17** | 2881.4 | 1110.0 | 1290.1 | 620.6 | 840.0 |
| Cluster 5 |  |  |  |  | **0.00** | 814.3 | 937.7 | 2021.0 | 4364.6 |
| Cluster 6 |  |  |  |  |  | **0.00** | 490.2 | **524.9** | **2382.2** |
| Cluster 7 |  |  |  |  |  |  | **452.10** | **808.45** | **1748.6** |
| Cluster 8 |  |  |  |  |  |  |  | **0.00** | **1587.2** |
| Cluster 9 |  |  |  |  |  |  |  |  | **0.00** |

**Supplementary Table 5.**
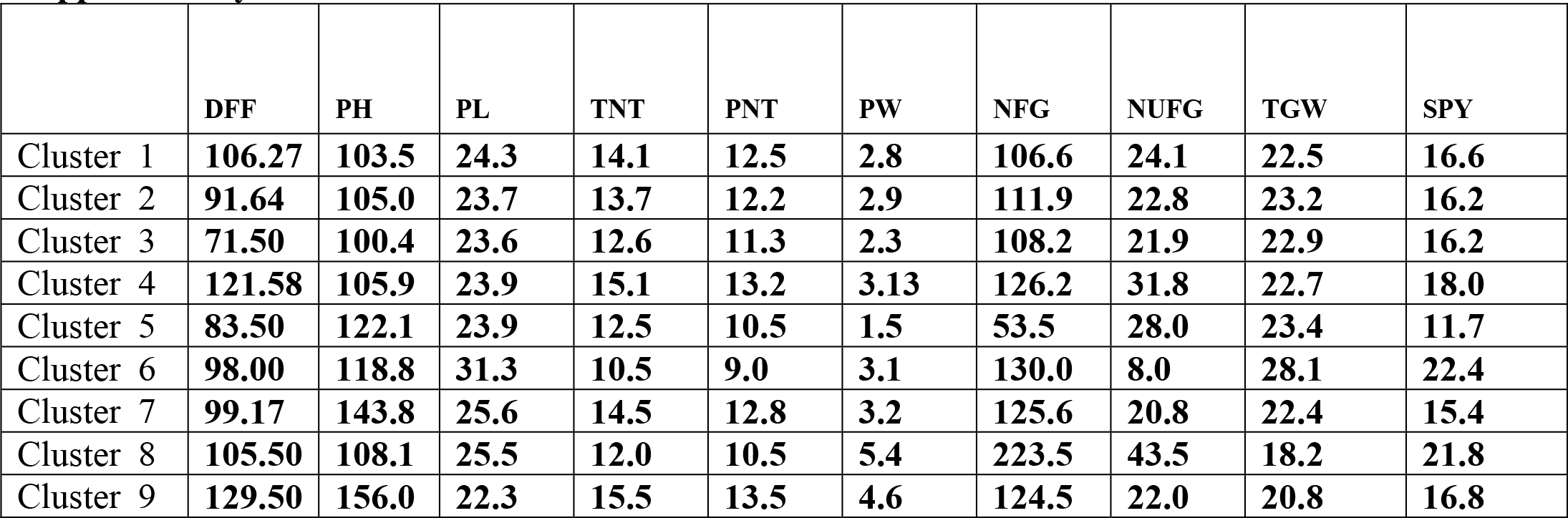
Cluster Means: Tocher Method.

**Supplementary Table 6.** Relative contribution of different characters to genetic diversity in germplasm lines.

| S.No | Characters | Contribution (%) |
| --- | --- | --- |
| 1 | DFF | 70.8 |
| 2 | PH | 16.16 |
| 3 | NFG | 4.73 |
| 4 | SPY | 3.71 |
| 5 | TGW | 3.56 |
| 6 | PW | 0.66 |
| 7 | NUFG | 0.15 |
| 8 | PL | 0.11 |
| 9 | PNT | 0.09 |
| 10 | TNT | 0.03 |

**Supplementary Table 7.** Direct and indirect effects of different characters on grain yield per plant in germplasm lines.

| Characters | DFF | PH | PL | TNT | PNT | PW | NFG | NUFG | TGW |
| --- | --- | --- | --- | --- | --- | --- | --- | --- | --- |
| DFF | **0.10** | 0.012 | 0.011 | 0.037 | 0.02 | 0.14 | 0.01 | 0.03 | -0.006 |
| PH | -0.019 | **-0.163** | -0.037 | 0.004 | 0.018 | -0.046 | -0.024 | 0.007 | -0.018 |
| PL | 0.022 | 0.043 | **0.193** | -0.043 | -0.058 | 0.020 | 0.023 | 0.012 | 0.028 |
| TNT | -0.337 | 0.025 | 0.205 | **-0.915** | -0.899 | 0.165 | 0.143 | -0.05 | 0.135 |
| PNT | 0.301 | -0.095 | -0.248 | 0.810 | **0.824** | -0.137 | -0.135 | 0.031 | -0.109 |
| PW | 0.024 | 0.028 | 0.010 | -0.018 | -0.016 | **0.099** | 0.068 | 0.025 | 0.033 |
| NFG | 0.015 | 0.015 | 0.012 | -0.016 | -0.016 | 0.070 | **0.102** | 0.025 | -0.010 |
| NUFG | 0.060 | -0.008 | 0.012 | 0.013 | 0.007 | 0.051 | 0.050 | **0.203** | 0.012 |
| TGW | -0.0006 | 0.0012 | 0.001 | -0.001 | -0.0014 | 0.003 | -0.001 | 0.0006 | **0.010** |
| SPY | 0.1682 | -0.142 | 0.163 | -0.129 | -0.104 | 0.251 | 0.243 | 0.276 | 0.076 |
| Partial R<sup>2</sup> | 0.0171 | 0.023 | 0.031 | 0.118 | -0.086 | 0.025 | 0.025 | 0.056 | 0.0008 |

**DFF- Days to 50% flowering, PH- Plant height, PL- Panicle length, TNT- Total number of tillers, PNT- Productive tillers,** **PW- Panicle weight, NFG- Number of filled grains, NUFG- Number of unfilled grains, TGW- Thousand grain weight, SPY-** **Single plant yield**

**Supplementary Fig.1.**
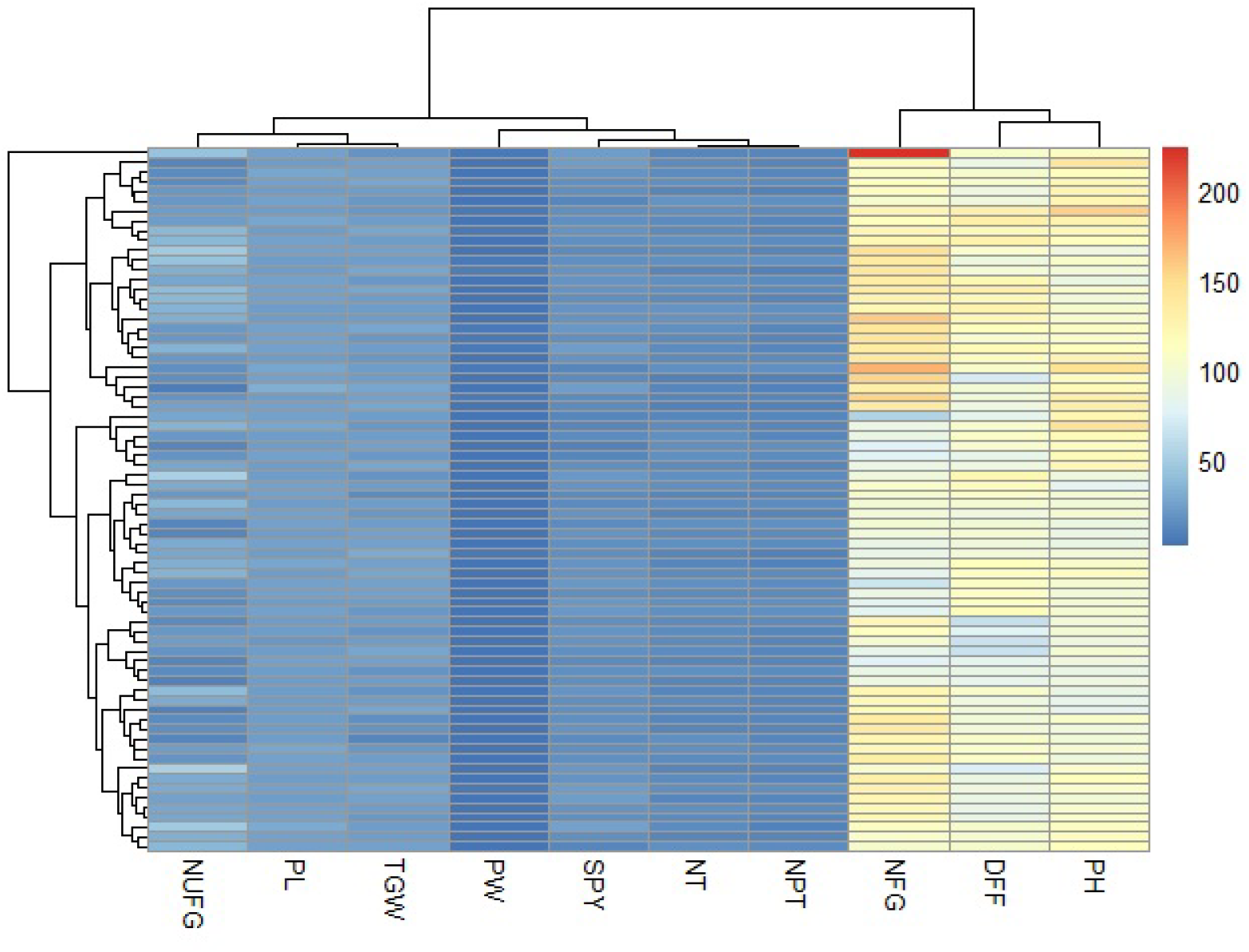
Heatmap depicting the genetic variability in germplasm lines for yield and its attributing traits.

**Supplementary Fig. 2.**
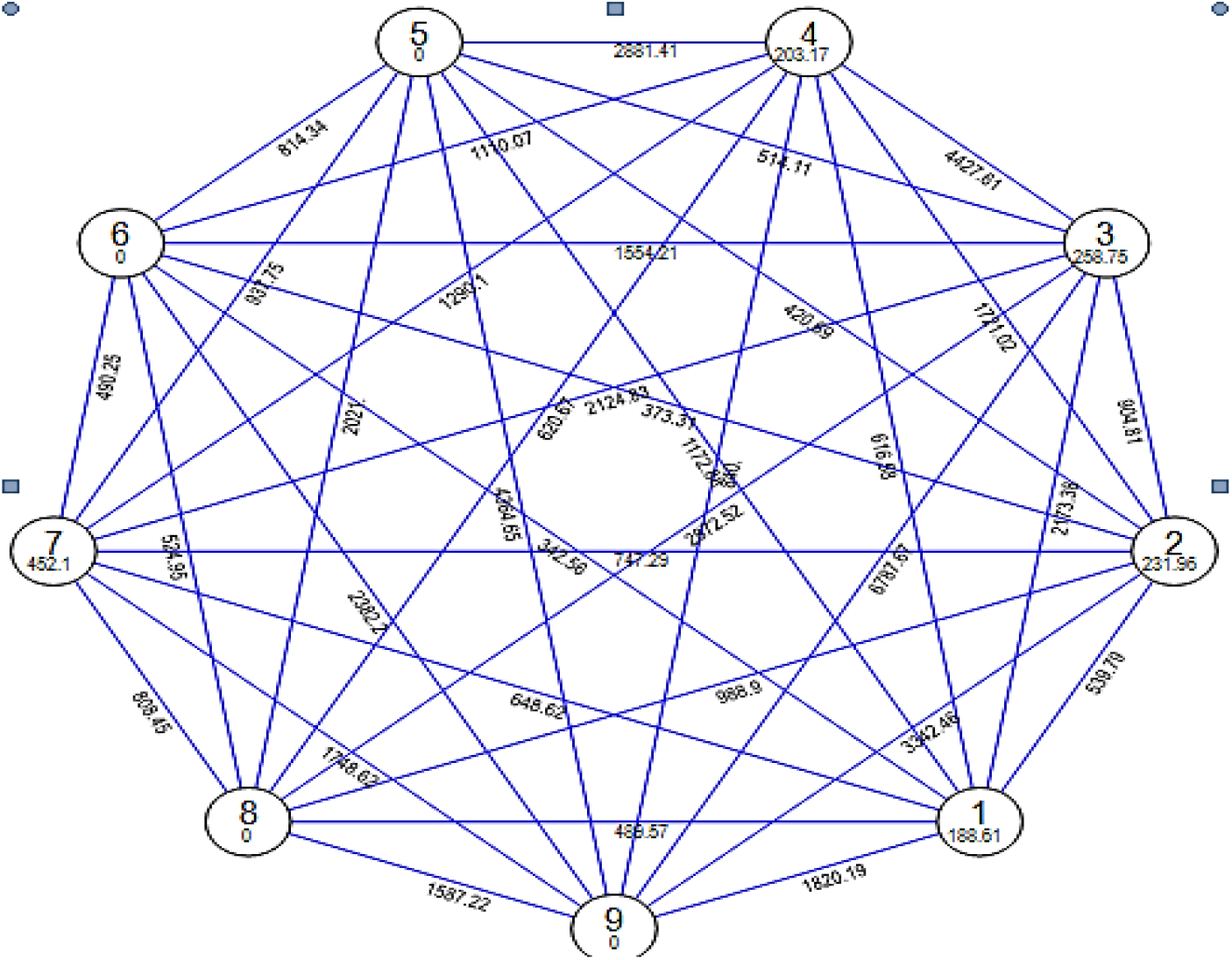
Cluster diagram of 72 germplasm lines based on D^2^ values by Tocher method.

